# Metabolic response to Parkinson’s disease recapitulated by the haploinsufficient phenotype of diploid yeast cells hemizygous for the adrenodoxin reductase gene

**DOI:** 10.1101/641621

**Authors:** Duygu Dikicioglu, James W. M. T. Coxon, Stephen G. Oliver

## Abstract

Adrenodoxin reductase, a widely conserved mitochondrial P450 protein, catalyses essential steps in steroid hormone biosynthesis and is highly expressed in the adrenal cortex. The yeast adrenodoxin reductase homolog, Arh1p, is involved in cytoplasmic and mitochondrial iron homeostasis and is required for activity of enzymes containing an Fe-S cluster. In this paper, we investigated the response of yeast to the loss of a single copy of *ARH1*, an oxidoreductase of the mitochondrial inner membrane, which is among the few mitochondrial proteins that is essential for viability in yeast. The phenotypic, transcriptional, proteomic, and metabolic landscape indicated that *Saccharomyces cerevisiae* successfully adapted to this loss, displaying an apparently dosage-insensitive cellular response. However, a considered investigation of transcriptional regulation in *ARH1*-impaired yeast highlighted that a significant hierarchical reorganisation occurred, involving the iron assimilation and tyrosine biosynthetic processes. The interconnected roles of the iron and tyrosine pathways, coupled with oxidative processes, are of interest beyond yeast since they are involved in dopaminergic neurodegeneration associated with Parkinson’s disease. The identification of similar responses in yeast suggest that this simple eukaryote could have potential as a model system for investigating the regulatory mechanisms leading to the initiation and progression of early disease responses in humans.

## Introduction

Iron is a crucial cofactor required for a number of essential cell functions for living organisms throughout the tree of life. It is a vital nutrient, which allows the transport of oxygen and the production of energy. Furthermore, it is essential for many metabolic and non-metabolic processes, including DNA repair and replication, as well as the regulation of gene transcription. These functions of iron are mainly based on its ability to donate electrons. Due to this property, iron may easily become a catalyst for reactions that facilitate the formation of free radicals. Therefore, iron is both essential for life and potentially toxic for the cell^1,2^. Numerous bacterial or fungal pathogens are highly dependent on iron supply and use different pathways to acquire iron from the environment or even steal it from their competitors or hosts^3,4^. Indeed, the battle between pathogenic microorganisms and host cells over iron has been proposed to play a critical role in infectious disease.

Many cell types or organisms lack effective means to secrete or excrete iron^5^. Due to the high toxicity of free iron, complex cellular regulatory mechanisms ensure its adequate acquisition, transportation, utilization, and elimination. This leads to the manifestation of an extremely fine-tuned metabolic system^1,2,6^. Disorders associated with both iron overload and deficiency relate to problems in this metabolic system^5,7^. Iron deficiency is the most prevalent nutritional abnormality in humans, although it has not been perceived as a life-threatening disorder. However, iron imbalance may lead to serious concerns in higher organisms such as alterations in circadian behaviour, and even neurodegeneration and cognitive impairment^8^. Both iron deficiency and excess was shown to lead to increased oxidative stress, and this often involves mitochondrial dysfunction, as observed in aging and associated degenerative diseases^9^. A major fraction of the oxygen taken up by eukaryotic cells is used by the mitochondria, which in turn, produce a substantial amount of cellular superoxide, and accumulate iron for the production of haem and iron-sulphur clusters^7^.

Iron-sulphur proteins are found in all life forms including archaea, bacteria and eukaryotes. Iron-sulphur clusters, polynuclear combinations of iron and sulphur atoms, are both structurally and functionally versatile, and are found in the most ancient components of living matter^10,11^ They can access various redox states, which makes them ideal for electron-transfer and redox reactions. Their structural, chemical, and electronic flexibility allow proteins that carry these clusters to participate in numerous biological functions, including electron transfer, substrate binding/activation, iron/sulphur storage, regulation of gene expression, and enzymatic activity^12^. All iron-sulphur clusters, regardless of whether they are utilised by mitochondrial, cytoplasmic, or nuclear proteins, are assembled in the mitochondria. Mitochondria play a key role in iron supply and utilisation in eukaryotic systems involving processes that are regulated and coordinated by iron-sulphur protein biogenesis^13^. These roles of mitochondria regarding iron metabolism, and particularly the generation of iron-sulphur clusters, have been conserved across eukaryotes from yeast to humans^14^.

Adrenodoxin reductase is a mitochondrial P450 flavoprotein, which catalyses essential steps in steroid hormone biosynthesis^15,16^. The enzyme is able to reduce a 2Fe-2S cluster protein, adrenodoxin, and was reported to be present in most metazoans and prokaryotes^17^. *ARH1* encodes the yeast adrenodoxin reductase homolog, which is an oxidoreductase of the mitochondrial inner membrane. The protein is involved in cytoplasmic and mitochondrial iron homeostasis, and plays a role in iron-sulphur cluster formation. It is one of the few mitochondrial proteins that are essential for viability in yeast^18–20^.

In our recent work, we investigated how yeast responded metabolically to an impairment in the function of the oxidoreductase encoded by *ARH1*. We showed that despite its essentiality, the yeast cells were able to cope with the loss of one copy of the gene (in an *ARH1/arh1Δ* hemizygote) without displaying any significant difference in growth rate, macronutrient utilisation, energy-associated product formation, or the sub-cellular accumulation of iron-species and copper characteristics. In order to benchmark our findings, we also investigated the response of yeast cells to the loss of a copy of *YFH1*, encoding the yeast frataxin homolog, which has a very similar function to *ARH1* in iron-sulphur cluster assembly, and also of *ATM1*, encoding the ABC transporter that exports mitochondrially synthesized precursors of iron-sulphur clusters to the cytosol. Our results showed that neither a *YFH1/yfh1Δ* nor an *ATM1/atm1Δ* hemizygote showed any significant difference form their cognate homozygous wild types in their metabolic or phenotypic responses^21^.

The assembly and functioning of iron-sulphur proteins that are employed outside of the mitochondria were reported to explain the indispensable nature of iron-sulphur cluster biogenesis for cell viability in virtually all eukaryotes. None of the mitochondrial iron-sulphur proteins in the yeast *Saccharomyces cerevisiae*, for example, is essential for life^13^. Most non-mitochondrial iron-sulphur proteins are involved in non-metabolic functions in the cell, and are therefore less likely to affect metabolic or phenotypic responses directly.

In this work, we attempted to identify the non-metabolic response to the impairment of iron-sulphur biogenesis. We conducted this analysis at the transcriptional level, but also made use of existing proteomics data to further extend the analysis. The collection of diploid yeast strains hemizygous for an individual gene (a hemizygote is a diploid strain in which one of the two copies of a given gene has been deleted) has proved invaluable in revealing both gene and drug function^22,23^. Here, we report that yeast hemizygous for the gene encoding the ortholog of the human adrenodoxin reductase enzyme substantially rewires its transcriptional regulation; a response not observed when a single copy of either of the other iron-sulphur cluster genes is deleted.

## Experimental

### Strains, cultivation conditions, and harvesting

Heterozygous deletion mutants *HO/hoΔ*, *ARH1/arh1Δ*, *ATM1/atm1Δ* and *YFH1/yfh1Δ* of *S. cerevisiae* strain BY4743 (background: *MATa/α his3Δ1/his3Δ1 leu2Δ0/leu2Δ0 lys2Δ0/LYS2 MET15/met15Δ0 ura3Δ0/ura3Δ0*) were used in this study, and the single-copy deletions were verified by PCR^24^. Qiagen DNeasy Blood & Tissue Kit was used for isolation and purification of DNA from the cell extracts as described in the manufacturer’s protocol.

Three different yeast strains were grown to an OD_600_ of 0.62-0.83, ensuring all cultures were in their exponential growth phase, at 30°C in YPD medium, allowing sufficient aeration in vented-cap Erlenmeyer flasks with shaking (200 rpm). Sample collection was adjusted by OD_600_ to harvest roughly 5×10^7^ cells per cryovial sample. The cells were flash-frozen in liquid nitrogen, and then stored at −80°C until further processing.

### RNA processing and microarray analysis

RNA extraction on the frozen samples was carried out using the Qiagen RNeasy Mini Kit, following the protocol for purification of total RNA from yeast. Cells were lysed mechanically using acid-washed beads and agitation in a FastPrep-24 5G Homogenizer (MP Biomedicals). The samples were agitated for 6×30s with 1 min resting on ice between lysis intervals. Samples with A_260_/A_230_ ratios < 1.8 were processed using the Qiagen RNeasy MinElute Cleanup Kit to reduce presence of contaminants.

Micro-volume UV-vis spectrophotometry was used for RNA quantification and purity (A_260_/A_280_) evaluation (NanoDrop ND-3000, Thermo Fisher Scientific Inc., U.S.A.). A microfluidics-based platform was used to check RNA integrity (Bioanalyzer 2200, RNA6000 Nanokit, Agilent Technologies, U.S.A.).

Microarray analysis was conducted as described in the GeneChip^®^ Expression Analysis Technical Manual (relevant kits, chips, and instrumentation: Affymetrix Inc., U.S.A.). Briefly, first-strand cDNA was synthesized from ca.100⍰ng of total RNA, and converted into ds DNA (GeneChip^®^ 3’ IVT Express Kit). Biotin-labelled aRNA was synthesised, purified and fragmented, and 5⍰μg of aRNA was loaded onto 169 format Yeast 2.0 arrays. Following hybridisation, the chips were washed and stained in the GeneChip^®^ Fluidics Station. The cartridges were stored at 4 C on ice and were scanned within 24 hours of preparation on the GeneChip^®^ Scanner 3000. The image files were processed and normalised with their quality assessed by dChip software^25^. MIAME^26^ compliant raw and processed files can be accessed from EBI’s ArrayExpress database^27^ (Accession no: E-MTAB-7648).

### Data analysis

Raw data were pre-processed prior to analysis. The dChip package was used to normalize probe intensity across the 12 samples to the lowest P-call percentage, and also to remove the background mismatch rate of probe hybridization, reducing the prevalence of false positives in the samples^25^. RMAexpress^28^ was used for background adjustment and quantile normalization of the data before transforming the probe intensities into log_2_ values. The normalised dataset was evaluated by a linear model that used median polish method^29^.

When the student’s t-test was used for statistical analysis, the Benjamini-Hochberg test was used to control the False Discovery Rate (FDR) at q < 0.05. One-way ANOVA was conducted at a significance threshold of 0.05. Significance Analysis of Microarrays (SAM)^30,31^ was employed as the test statistic for the statistical evaluation of gene expression with 90^th^ percentile False Discovery Rate and 2-fold expression change as additional constraints on evaluation.

Latent class analysis and correspondence analysis was carried out as described by Kamakura and Wedel^32^. Correspondence analysis was used to linearly transform the transcription factor (TF)-target gene associations resulting in a discrete data matrix. The threshold for discrimination was |±0.15| from the origin. Latent class analysis was used to classify categorical data. The classification was carried out using two approaches: the corrected-Akaike’s information criterion (CAIC)^33^ and log-likelihood ratio test^34^. The number of classes was selected based on domain-usefulness such that the highest number yielding all classes to be non-small is selected, where small size is described as containing less than 5% of the total sample group, i.e. the gene subset, in this case. This rule, which has long been used in practice as a part of the idea of domain-usefulness, has recently been discovered also to have some theoretical justification^35^.

Princeton GO tools were employed for Gene Ontology Enrichment Analysis accessed on 07/2018^36^. The background list was 7166 transcripts. The significant ontology terms listed had a Bonferroni-corrected p-value of less than 0.01. False Discovery Rates were also calculated and those terms with q-value > 0.05 were excluded from further analysis. Protein expression data used for comparison was accessed from Paulo et al. 2016^37^. The yeast mRNA decay data was accessed from Munchel et al. 2011^38^. Spearman correlation (implemented in MATLAB) was used to evaluate the relationship between mRNA decay and protein expression. Transcriptional regulatory network interactions were accessed from YEASTRACT^39^ (access date 07/2018). The BLASTP Tool^40^ was used via UniPROT^41^ for the identification of sequence similarities. Human protein homology for the yeast proteins was accessed from Saccharomyces Genome Database^42^, and the genetic interactions for yeast were accessed from TheCellMap^43^ on 22/06/2018.

## Results and discussion

### Transcriptional and proteomic response of yeast to impairment in iron-sulphur cluster (ISC) machinery

The transcriptional response of diploid yeast to the loss of a single copy of either *ARH1*, *ATM1* or *YFH1* was investigated in comparison to that of a control strain (ESI). Haploinsufficiency in yeast was shown to be primarily caused by insufficient protein production^44^, and in the budding yeast *S. cerevisiae* protein abundance was shown to match gene copy number quantitatively^45^, demonstrating an inability to compensate for reduced gene dosage. All three genes under investigation in this work have key roles in the biogenesis of iron-sulphur clusters in the mitochondria, and their null mutants were reported to be inviable in large-scale genetic surveys^42^.

Following a preliminary statistical analysis of the transcriptome data (ESI2), a supervised learning algorithm that relies on non-parametric statistics was selected as a suitable tool for mining this genome-wide expression dataset. The expression levels of 69 genes were significantly different in *ARH1/arh1Δ* at a confidence level of 90%, accompanied by a 2-fold change in expression. A significant enrichment was identified among those genes for iron ion transmembrane transporter activity (p-value = 0.005). In contrast, the expression levels of only 40 and 5 genes were different in *ATM1/atm1Δ* and *YFH1/yfh1Δ*, respectively. The *YFH1/yfh1Δ* subset was not significantly enriched for any particular biological processes or molecular functions. This could indicate that the yeast could successfully cope with the reduced gene dosage for *YFH1*; although an inviability of a null mutant would indicate that complete loss of functionality would not be tolerable. The genes that displayed a significant change in their expression in the *ATM1*-impaired yeast were significantly associated with ribosome biogenesis (p-value = 1.68 × 10^−7^), and were significantly involved in DNA-directed 5’-3’ RNA polymerase activity (p-value = 0.002) (Fig.1, ESI).

**Fig. 1.**
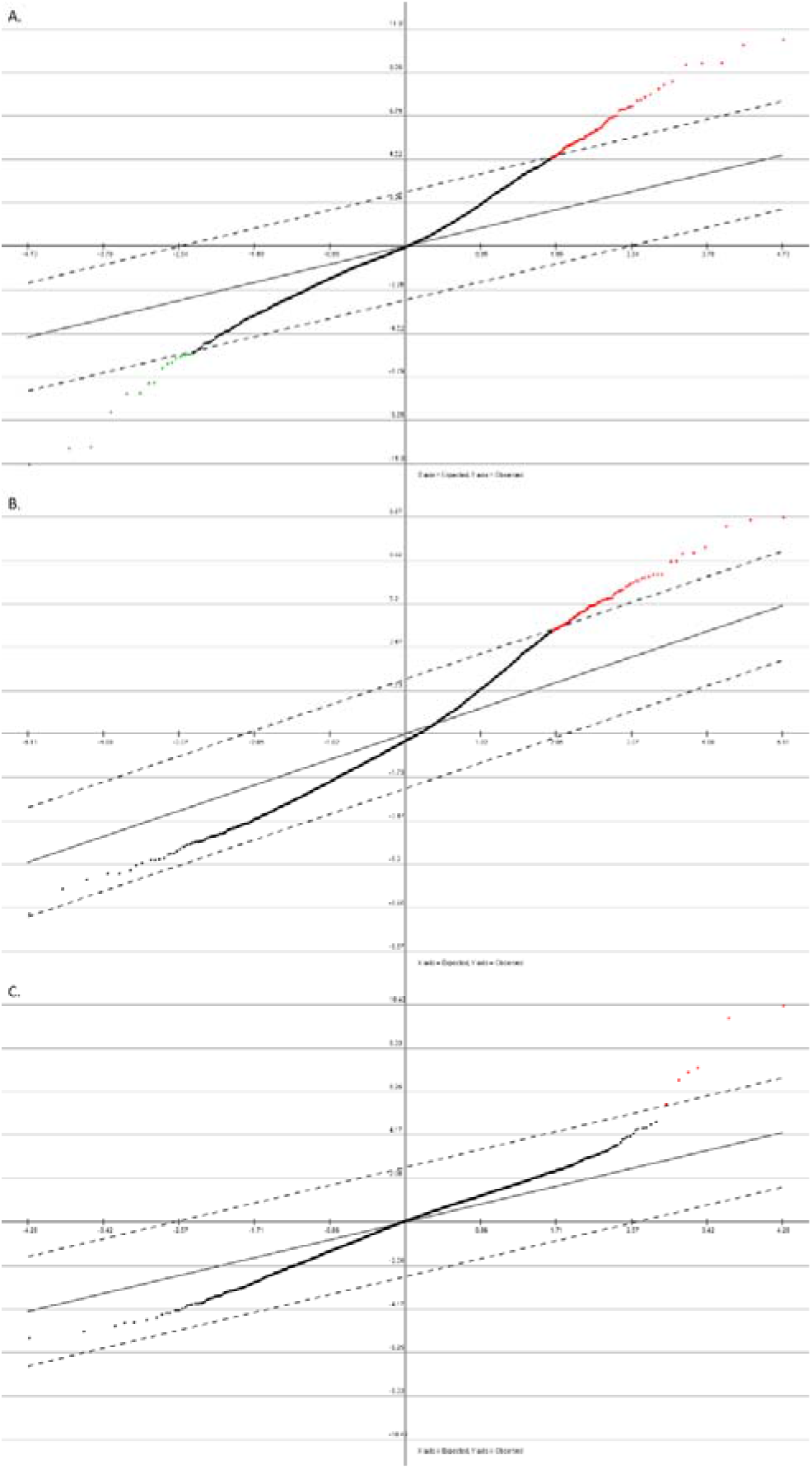
Significance analysis of the gene expression by SAM for *ARH1/arh1Δ* (A), *ATM1/atm1Δ* (B), and for *YFH1/yfh1Δ* (C). The expected relative expression values are provided on the abscissa and the observed relative expression values are given on the ordinate. Note all expression values are log_2_ converted. The solid line denotes expected value = observed value, and the dashed lines denote the 90% confidence intervals after FDR correction. Positive significant genes are labelled in red and negative significant genes are in green, to provide a summary view of the fraction of genes whose expression was significantly different in these mutants.

Protein levels during growth on glucose were available for 87% of the genes, whose gene expression levels were measured in our study. The protein levels were available for approximately the same fraction of the genes (89%) that were differentially expressed in response to the loss of a single copy of the iron-sulphur cluster (ISC) genes, allowing a consistent analysis to be made. However, there was no significant correlation between these protein levels and the transcript levels of the subset of differentially expressed genes (even without the 2-fold change in expression constraint) in these mutants (ρ = 0.11, p- value = 0.24 for *ARH1/arh1Δ*, and ρ = 0.33, p- value = 3.08 × 10^−4^ for *ATM1/atm1Δ*, and not applicable for *YFH1/yfh1Δ* due to unavailability of protein abundance data). This lack of correlation between the transcript and protein levels, not uncommon in yeast^46^, led us to investigate post-transcriptional modification events in yeast, mainly that of mRNA turnover. However, no positive or negative correlation was observed between protein abundance and mRNA half-lives (ρ = −0.37, p- value = 5.76 × 10^−5^ for *ARH1/arh1Δ*, and ρ = −0.33, p- value = 3. 82 × 10^−4^ for *ATM1/atm1Δ*, and not applicable for *YFH1/yfh1Δ* due to unavailability of protein abundance data for the proteins encoded by the genes in that subset).

### Role of the transcriptional regulation in the impairment of the ISC machinery

The relationship between transcription factors (TFs) and their target genes was investigated to explore the rewiring of yeast cellular networks in response to the loss of a single copy of a gene encoding a component of the ISC machinery. Only two genes whose expression levels significantly changed in response to the impairment in *ARH1* encoded TFs: *IME1*, and *MGA1*; whereas no such genes were identified in *ATM1/atm1Δ* and *YFH1/yfh1Δ*. Furthermore, the protein abundance levels for Ime1p and Mga1p were not captured in the proteomics dataset used here. Therefore, shared transcriptional regulatory patterns were identified instead for the 134-, 132-, or 6-gene subsets that were transcriptionally responsive (significant, but without fold-change considerations) to the heterozygosity of *ARH1*, *ATM1*, or *YFH1*, respectively. These patterns were determined to be similar for the *ARH1/arh1Δ* and *ATM1/atm1Δ* subsets (ESI). These gene subsets and transcription profiles were then analysed using various TF-target gene association measures (see ESI2 for details).

In light of this inference, the TFs whose target gene expression profiles varied significantly in response to the heterozygosis of *ATM1* or *ARH1* were investigated solely from a functional point of view. The subset of genes identified in the *YFH1/yfh1Δ* comprised just six genes, making a systematic interpretation of the results problematic. Therefore, the differences and similarities observed between this subset and the others were excluded from further consideration (ESI).

The TFs, which had a high impact on the regulation of the expression of the genes in the two subsets, were identified and functionally investigated for their potential effect on the observed transcriptional response. Only 7 and 8 TFs (which we shall term ‘major controllers’) were reported to regulate the expression of more than 50% of the genes whose transcript levels changed significantly in *ARH1/arh1Δ* and *ATM1/atm1Δ* strains compared to the wild-type diploid. Six of these TFs (Ace2p, Ash1p, Cst6p, Gcn4p, Msn2p, and Rap1p) were shared between the two subsets, and these TFs were almost exclusively involved in regulating the transcription related to the cell cycle and division^42^.

On average, each major controller TF had a documented association with more than 15% of the genes in each subset (Fig.S1B in ESI2). However, more than 25% of all yeast TFs were reported to regulate the expression of a greater fraction (than 15%) of the genes in their respective subsets as verified by both binding and gene expression data. This populated a large TF pool for either mutant; the pool comprised 51 TFs for each of *ARH1/arh1Δ* and *ATM1/atm1Δ*, of which 43 were shared. Despite the direct involvement of *ARH1* and *ATM1* in iron-related processes, only three TFs were identified either to regulate iron metabolism, or to require iron-containing cofactors; Aft1p, Hap2p, and Hap4p in *ARH1/arh1Δ*, and Aft1p, Cad1p, and Hap2p in *ATM1/atm1Δ*. This limited inference of iron-linked regulation by transcription factors was in line with the minimal response observed in the expression of iron-associated genes in the transcriptome analysis. On the other hand, 27% and 24% of these TFs were responsive to oxygen availability and oxidative stress in *ARH1/arh1Δ* and *ATM1/atm1Δ*, respectively.

### Categorical classification to identify the gene regulatory relationships

The complete dataset was next employed in classification analysis, which is a form of clustering for categorical data as identified in these TF-target gene association data (ESI, ESI2). The subset of genes that displayed a significant change in response to the loss of a single copy of *ATM1* or *ARH1* were then grouped into 5 and 7 mutually exclusive and exhaustive classes, respectively, based on their pattern of transcriptional regulatory responses determined by all yeast TFs acting as categorical variables in Latent Class Analysis.

All classes in this analysis for the *ATM1/atm1Δ* strain were significantly associated with ribosomal and translation-associated processes (p-value = 0.01), and thus did not contribute further to our initial analysis solely conducted on the basis of gene transcription patterns (see ESI). Furthermore, the TFs targeting these genes were predominantly responsive to oxidative stress (see previous section for a detailed discussion). This relationship between ribosomal events and oxidative stress agrees with early reports documenting ribosomal rRNA damage to be elevated by an increase in the intracellular level of reactive oxygen species^47^.

In contrast the classes identified for the *ARH1/arh1Δ* yeast were discretely and significantly associated with a variety of different and seemingly unrelated processes (p-value = 0.01). One class was populated with genes involved in iron assimilation (p-value = 0.005), and another class in tyrosine biosynthesis (p-value = 0.007). Alterations involving iron-associated processes in response to the hemizygosity of *ARH1* were not unexpected and were consistent with our initial review of the gene expression data. However, the involvement of tyrosine biosynthesis was a novel finding elucidated by latent class analysis.

Another class identified by this analysis was enriched for genes involved in oxidoreductase activity (p-value = 0.007), and yet another class included two of the three genes constituting the Rix1-complex; *IPI1* and *IPI3* (p-value = 0.0007). Members of the RNA-processing Rix1-complex were previously reported to be responsive to oxygen levels, particularly to hypoxia^42,48^. The highly conserved members of this complex were also haploinsufficient in the presence of chemical-induced oxidative stress^49^. Thus, latent class analysis identified oxidation-related gene targets in two different classes, which were possibly associated with the oxidation-related regulatory activity identified in the aforementioned TF analyses.

### Iron and tyrosine metabolism in response to reduced adrenodoxin levels and its potential implications for neurodegenerative disorders

In order to investigate the link between iron assimilation and tyrosine biosynthesis in conjunction with oxidative stress, we constructed a protein interaction network in yeast. The first neighbour interactors of the proteins involved in iron assimilation (Ftr1p, Fet3p, Fet5p, and Gmc1p) and tyrosine biosynthesis (Aro7p, Aro8p, Aro9p, and Tyr1p) were identified and the crosstalk between the two networks was determined. Among the shared first-neighbour interactors, only one protein was known to be associated with oxidative stress in yeast, Hsp104p. This heat shock protein has been reported to function as a disaggregase, that assists the refolding and reactivation of previously denatured, aggregated proteins^42^ (Fig.2).

**Fig. 2.**
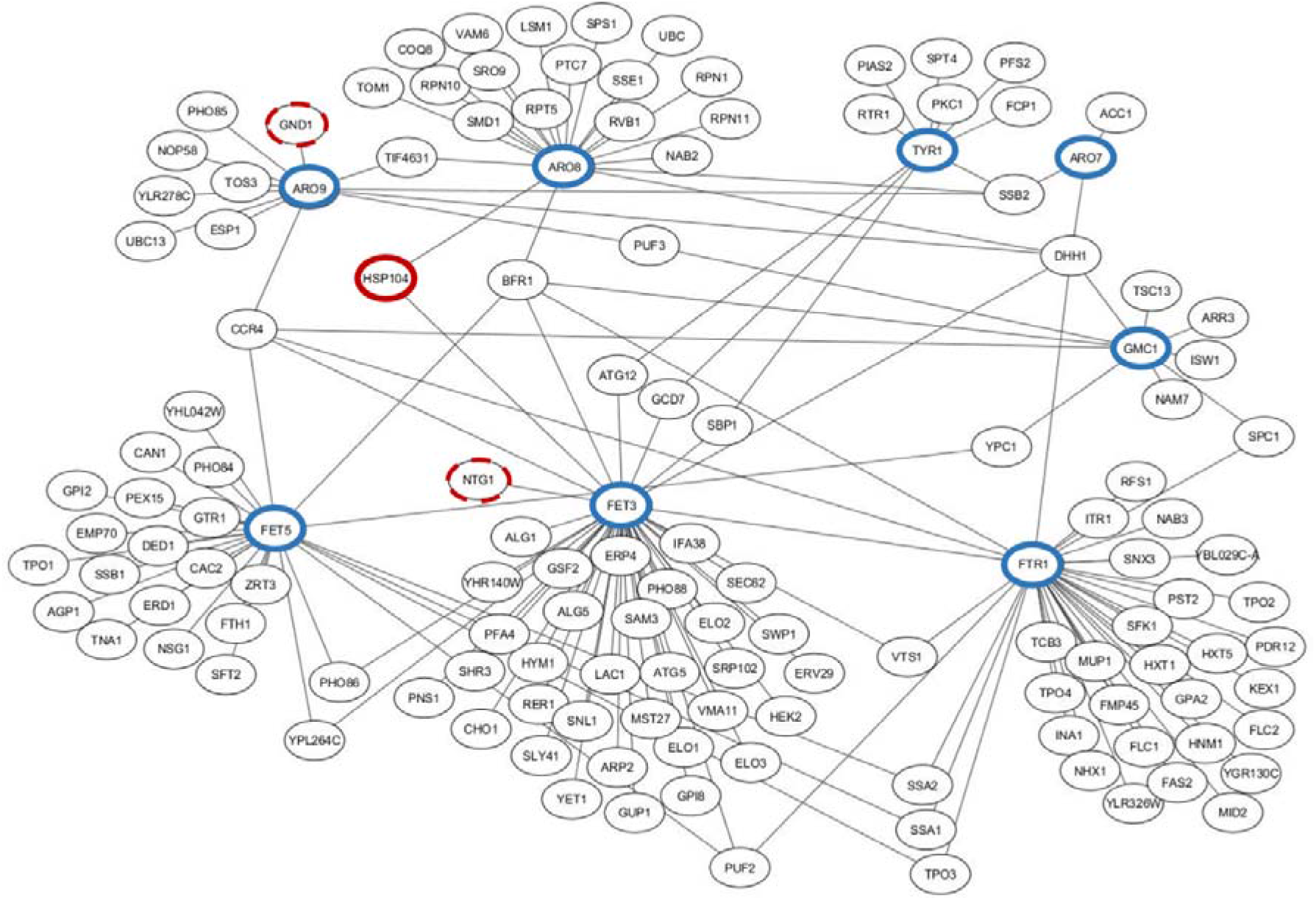
Crosstalk between iron assimilation and tyrosine biosynthesis in yeast. The protein-protein interaction network was constructed between the genes encoding the proteins involved in iron assimilation and tyrosine biosynthesis, encircled in blue colour. The first neighbours with a role implicated in oxidative stress are encircled in red. The dashed line indicates that the first neighbour does not establish a cross talk between the processes while the solid line indicates the gene encoding the cross-talk protein.

None of the genes encoding the iron assimilation or tyrosine biosynthesis proteins that were identified in this study are homologous to human genes that were known to be implicated in Parkinson’s disease. However, some functional associations exist. An important disease marker, alpha-synuclein, was shown to inhibit the retrograde recycling of the yeast Fet3p/Ftr1p heterodimer at low external iron concentrations^50^. Aro8p and Aro9p were recently identified as the major kynurenine aminotransferases in yeast, catalysing the deamination of kynurenine to kynurenic acid^51^. Kynurenic acid was reported to have a role in interfering with dopaminergic neurotransmission, highlighting its relevance for Parkinson’s disease^52^. The disorder was reported to be accompanied by abnormalities in the kynurenine pathway with extended associations implicated by defects in respiratory complex I in the mitochondria^53,54^; the pathway intermediates were even suggested as plasma biomarker candidates for the disease^55^. Furthermore, iron and tyrosine are two metabolites that have been strongly linked with Parkinson’s disease. Parkinson’s patients preferentially lose the dopamine-producing neurons in their midbrain, and consequently have low dopamine availability^56^. Patients with the disorder also have reduced levels of norepinephrine^57^; L-DOPA supplementation, a standard treatment for Parkinson’s patients, lowers the norepinephrine level even further. Tyrosine is converted to L-DOPA; the precursor for the neurotransmitter, dopamine through the catalytic action of the iron-binding tyrosine hydroxylase^58^. Dopamine, in turn, acts as the main precursor for norepinephrine^59^. Dietary supplements of amino acids, including tyrosine, have been given to Parkinson’s patients to address the complications caused by treatment with L-DOPA^60^.

A characteristic feature of the Parkinson’s brain is the substantial accumulation of iron. Although it is not yet understood clearly whether iron accumulation is the cause or the consequence of the disorder, there is evidence that the neurodegeneration could be initiated by the potent redox couple formed by iron and dopamine itself^61^, leading to substantial oxidative stress. Whether this is the case or not, it is clear that our yeast model replicates the metabolic fingerprint of this neurodegenerative disorder in this genetically and physiologically malleable simple eukaryote.

The re-wiring of the regulation of gene transcription in the *ARH1/arh1Δ* yeast hemizygote mirrors the metabolic response exhibited in Parkinson’s disease. On a genetic basis, the role of the tyrosine pathway was particularly interesting in this context. Although mitochondrial iron metabolism^14^ and many different types of oxidative stress response^62,63^ are highly conserved across species, yeast lacks the tyrosine hydroxylase enzyme, which would catalyse the first step in the synthesis of catecholamines including dopamine and norepinephrine. The human tyrosine hydroxylase shares *ca*. 35% similarity with two proteins of the closely related yeast, *Zygosaccharomyces rouxii*. These have more than 20% sequence similarity to a number of *S. cerevisiae* proteins, of which Flo9p, Flo11p, and Hkr1p are particularly interesting due to their role in calcium-associated and aggregation-related processes^42^ (see ESI2 for detailed analysis).

## Conclusions

In this work, we investigated the transcriptional and gene regulatory response of diploid yeast to the loss of a single copy of one of the genes that play essential roles in the iron-sulphur cluster machinery: *ARH1*, *ATM1*, or *YFH1*. The transcriptional response and the governing gene regulatory events were very limited for the *YFH1/yfh1Δ* mutant, indicating that yeast could cope with the loss of a single copy of this gene relatively successfully. A modest response, coupling ribosomal processes with oxidative stress, was observed for the *ATM1/atm1Δ* cells. This novel association, uncovered by latent class analysis, was the only significant response of diploid yeast to the loss of a single copy of *ATM1*. In contrast, the response of diploid yeast to the deletion of a single copy of *ARH1* involved a greater range of biological processes than was the case for the other two hemizygous mutants, and included tyrosine biosynthesis, iron assimilation, and the oxidative stress response. These three processes are substantial contributors to the cellular phenotype of Parkinson’s disease and this suggests that the yeast *ARH1/arh1Δ* hemizygote recapitulate the disease phenotype. Although it is not yet clear how, and to what extent, haploinsufficiency is known to be involved in human hereditary disease^44^. Furthermore, the yeast *Saccharomyces cerevisiae* has long been employed as a model system for both understanding cellular mechanisms in, and discovering potential drugs for treating, Parkinson’s disease^64–67^. Therefore, our findings emphasise the potential for exploiting yeast, specifically those mechanisms involving adrenodoxin reductase homolog, as a model cellular system for investigating the onset and early progression of Parkinson’s disease.

## Supporting information

ESI

ESI2

## Conflicts of interest

There are no conflicts to declare.

## Acknowledgements

We thank Andy Hesketh for discussions on microarray analysis, Mark Kristiansen and Kerra Pearce from the Genomics Centre at University College London Great Ormond Street Institute of Child Health for the provision of access to Affymetrix Scanners. We also thank Gabriele S Kaminski Schierle for extensive discussions on Parkinson’s disease and the role of iron and tyrosine. The authors gratefully acknowledge the funding from the Leverhulme Trust and the Isaac Newton Trust (ECF-2016-681 to DD), EC 7^th^ FP (BIOLEDGE Contract no: 289126 to SGO), BBSRC (BRIC2.2 to SGO). None of the funding agencies had a role in study design, data collection or interpretation, or the decision to submit the work for publication. The authors declare no competing interests.

