## Supplementary material for "Metabolic response to Parkinson’s disease recapitulated by the haploinsufficient phenotype of diploid yeast cells hemizygous for the adrenodoxin reductase gene": ESI2

**Electronic Supplementary Results**

**Preliminary statistical analysis on the transcriptome data**

A simple analysis of significance via the Student’s t-test followed by FDR correction identified only 20 transcripts, whose expression significantly differed between *ARH1/arh1Δ* and *HO/hoΔ*. This group of genes were not significantly associated with any Biological Process, Molecular Function or Cellular Component Gene Ontology Terms (p-value < 0.05). The expression of genes did not display any significant difference in *ATM1/atm1Δ* and *YFH1/yfh1Δ* strains, in comparison to that of the control strain. Relaxing the significance threshold for evaluating differential expression to a p-value of 0.1 did increase the size of the differentially expressed gene set for *ARH1/arh1Δ* up to 774, but did not have any effect on those of the *ATM1*- or *YFH1*-impaired strains. This indicated that these strains were relatively more successful at complementing the loss of a single copy of these genes than the *ARH1/arh1Δ* could. One-way ANOVA analysis did not identify any genes, whose expression displayed a significant difference between these mutants and that of the control strain.

The overall nature of the dataset indicated that the distribution of the data points towards the extremes of gene expression (i.e. the tail was thinner than the tails of the normal distribution (kurtosis < 0.55, as opposed to 3 for normal distribution). This non-normality led to the implementation of alternative methods for the analysis of the significance of the changes in gene expression. A supervised learning strategy was thus selected for further analysis.

**TF-target gene association measures for *ARH1/arh1Δ* and *ATM1/atm1Δ* subsets**

The average “TF-binding measures” as well as the distribution profiles of these “measures” were determined for these subsets. The average fraction of TFs that can bind to each gene in the subset (15.92 ± 0.31%), the fraction of transcripts that possess binding sites to accommodate more TFs than the reported mean (44 ± 4.95%), and the average fraction of the subset of genes targeted by each yeast TF (15.68 ± 0.44%) were similar for the *ARH1/arh1Δ* and *ATM1/atm1Δ* subsets. Not only were these mean measurements, but also the frequency and the cumulative distribution profiles of these parameters across the complete range of values similar for these subsets (Fig. S1). Furthermore, the number of TFs that can bind to more than half of the genes constituting the subset (8 ± 0.71), or to more than the average number of genes targeted by each TF determined for each subset (44 ± 0.71) were also comparable (ESI).

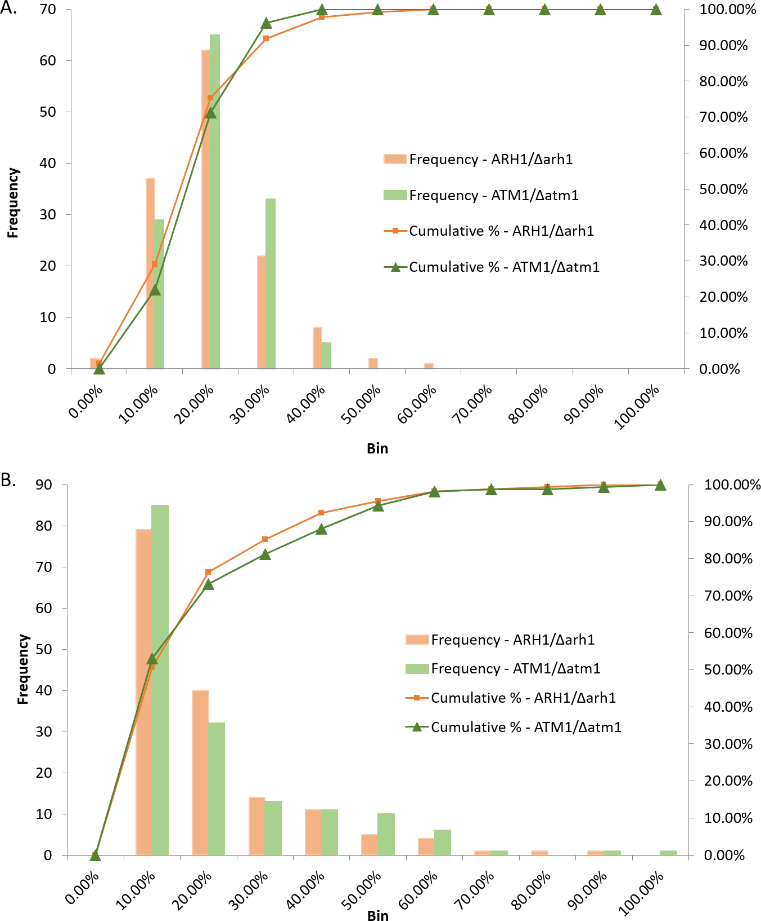

**Fig. S1** Frequency and cumulative distribution profiles of TF-binding measures for *ARH1/arh1Δ* and *ATM1/atm1Δ*. The distribution of the fraction of TFs that can bind to each gene whose expression significantly varied in their respective mutants is displayed in (A), and the distribution of the fraction of subset of transcripts with a significant change targeted by each yeast TF is displayed in (B). Shades of orange and green are used to represent data for *ARH1/arh1Δ* and *ATM1/atm1Δ*, respectively. The frequency of the observations are classified into ten bins (from 0% to 100%), and the cumulative percentages are calculated through the summation of consecutive bins starting from the first bin (0-10%) onwards.

The comparability of these two subsets with respect to TF-target gene association measures eliminated the possibility of any mechanistic or physical differences, which could become prominent merely due to numerical dissimilarities.

**Multivariate analysis to identify TF-target gene relationships**

A correspondence analysis model was constructed to assist the visualisation of the relationship between the pooled TFs and their differentially expressed target genes. The genes that were highly discriminated in this analysis or the TFs identified as the discriminating attributes did not share common features to classify them into a specific biological process. A remarkable number of latent factors were observed to have similar contributions in magnitude explaining the variance in the data. A two- or three-dimensional representation, which would be convenient for visualisation purposes, could only represent less than 25% of the total variance in the dataset, thus rendering the analysis insufficient for revealing any biological insight (see ESI).

**Functionally non-conserved proteins contributing to similar phenotypes in the Baker’s yeast and in human**

The human tyrosine hydroxylase (EC 1.14.16.2) has 36.8% identity with an uncharacterised protein (UniProt ID: A0A1Q3A628) and 29.9% identity with the **eukaryotic translation initiation factor 3 subunit A** from *Zygosaccharomyces* *rouxii*, which belongs to the Saccharomycetaceae family along with *S*. *cerevisiae*, and this is the nearest similarity in terms of sequence or functional conservation. Furthermore, A0A1Q3A628 protein has sequence similarity with a number of *S*. *cerevisiae* proteins (identity range: 23.1 - 29.2%, confidence e-value range:$1.3\times{10}^{-79}-4.0\times{10}^{-38}$). These are the cell wall proteins Dan4p, Sed1p, Fig2p, Fpg1p, and Awa1p; the GPI-anchored cell surface glycoprotein Flo11p, which is required for the formation of fibrous interconnections between cells of a wild strain; the lectin-like Flo9p, which is similar to Flo1p; and Hkr1p, the mucin family member osmosensor, which also functions in the filamentous growth pathway^1^. Flo9p and Flo11p play a role in Flo1-phenotype flocculation, which is calcium-dependent and constitutive^2^.

Although α-synuclein, the major contributor to the pathological hallmark of Parkinson’s disease, was not identified to bear sequence similarity to either Flo9p or Flo11p directly, it was associated with Flo1p, Flo5p, and Flo9p. All these proteins have a role in Flo1-phenotype flocculation and share sequence similarity with a flocculin (YFA0S_01e07294g) from the yeast *Cyberlindnera* *fabianii* (38.2% identity and e-value:2$.3\times{10}^{-4}$ human-yeast alignment, 28.2% identity and e-value:2$.5\times{10}^{-47}$ yeast-yeast alignment). These associations may imply a possible indirect relationship between yeast proteins with a function in agglomeration and key proteins implicated in the Parkinson’s disease pathology.
